# LC-MS system for automatically collecting time-resolved metabolomics data of cultured cells

**DOI:** 10.1101/2024.10.11.617934

**Authors:** Carly C.Y. Chan, Ryan A. Groves, Ian A. Lewis

## Abstract

Temporal metabolic dynamics are a critical, but difficult to study aspect of metabolism. To address this, we developed a liquid chromatography-mass spectrometry (LC-MS) system, temporal uptake and nutritional analysis (TUNA), to automatically collect time-resolved metabolomics data of cultured cells. TUNA enables sub-minute sequential sampling, has broad metabolite coverage, supports robust metabolite identification, can monitor over 72 conditions in parallel, and can be implemented in most LC-MS laboratories. We used TUNA to monitor temporal dynamics of uropathogens (*Escherichia coli* and *Proteus mirabilis*) and identify novel metabolic phenotypes that cannot be captured from a single time point.

## Main

Metabolomics captures a broad transect of the metabolic network, but has a critical blind spot in its ability to assess temporal shifts in metabolic phenotypes. This is problematic in the context of microbiology, where cultures can exhibit major shifts in metabolism resulting from changing nutrient availability and cell density. These metabolic shifts can be dramatic, in some cases leading to the reversal of fluxes.^1,2^ For example, *Escherichia coli* preferentially catabolize glucose and secrete succinate, but will subsequently catabolize succinate when nutrients become limiting.^1^ Likewise, *Clostridium acetobutylicum* produce acetate and butyrate under nutrient-rich conditions, but subsequently consume these molecules and convert them to acetone and butanol.^2^ In addition to these dramatic flux reversals, cells have hierarchical carbon utilization, wherein nutrients are preferentially catabolized in a specific sequential order.^1^ Capturing these temporal trends is challenging using conventional metabolomics methods.

The established strategy for monitoring temporal metabolic dynamics is to manually collect time series data of cell cultures^3,4^ (Supplementary Fig. 1). This approach is effective, but laborious, and introduces multiple sources of error. To address this, several automated monitoring technologies have been proposed. These platforms include biosensors^5^, microfluidics^6^, nuclear magnetic resonance (NMR)^7^, and mass spectrometry^8^. Each of these strategies can be harnessed for real-time metabolomics analysis, but the current solutions have limitations: Biosensors are generally limited to a handful of target molecules^5^; Microfluidics sampling require custom devices that are inaccessible to most laboratories^6^; NMR requires samples to be grown within NMR tubes, which have questionable physiological relevance^7^; Mass spectrometry strategies inject raw cell cultures directly into the instrument via flow injection^8^. Out of all these approaches, the liquid chromatography-mass spectrometry (LC-MS) system supports the broadest metabolite coverage. However, injecting unprocessed cultures into the instrument leads to metabolite extraction and deposition of cell material within the ion optics. Moreover, this approach is not compatible with chromatography, which creates sensitivity problems from ion suppression and prevents robust LC-MS metabolite assignment. Currently, there are no strategies for automatically collecting physiologically relevant metabolomics data over time that provide broad metabolite coverage, unambiguous metabolite identification, can be conducted over long time courses, and can be implemented in routine laboratory settings.

To address limitations, we developed an automated temporal uptake and nutritional analysis (TUNA) system for cultured cells. Four recent advances in our group laid the foundation for TUNA (Fig. 1). Firstly, we developed a high-throughput LC-MS method for profiling hydrophilic molecules present in culture media.^4,9^ Secondly, we developed a microbial containment device (MCD) that separates the cell culture well from the sterile analytical well via a semi-permeable 0.2 μm filter.^10^ This allows cell-free metabolites to be sampled over time and injected directly into the LC-MS without introducing microbial contaminants into the system^10^, thereby permitting the use of chromatography to minimize ion suppression and enable robust metabolite identification. Thirdly, we developed a biomarker enrichment medium (BEM), a chemically defined growth medium engineered specifically for LC-MS analysis of microbial metabolism.^11^ Lastly, we developed a boundary flux analysis (BFA) strategy for characterizing the phenotypes of cells based the rates with which metabolites are either consumed or secreted in growth media.^9,12^ TUNA leverages each of these components as well as one novel element: we use the LC-MS sample changer as a microbial incubator, which can be calibrated to 37 °C (Fig. 1).

**Figure 1.**
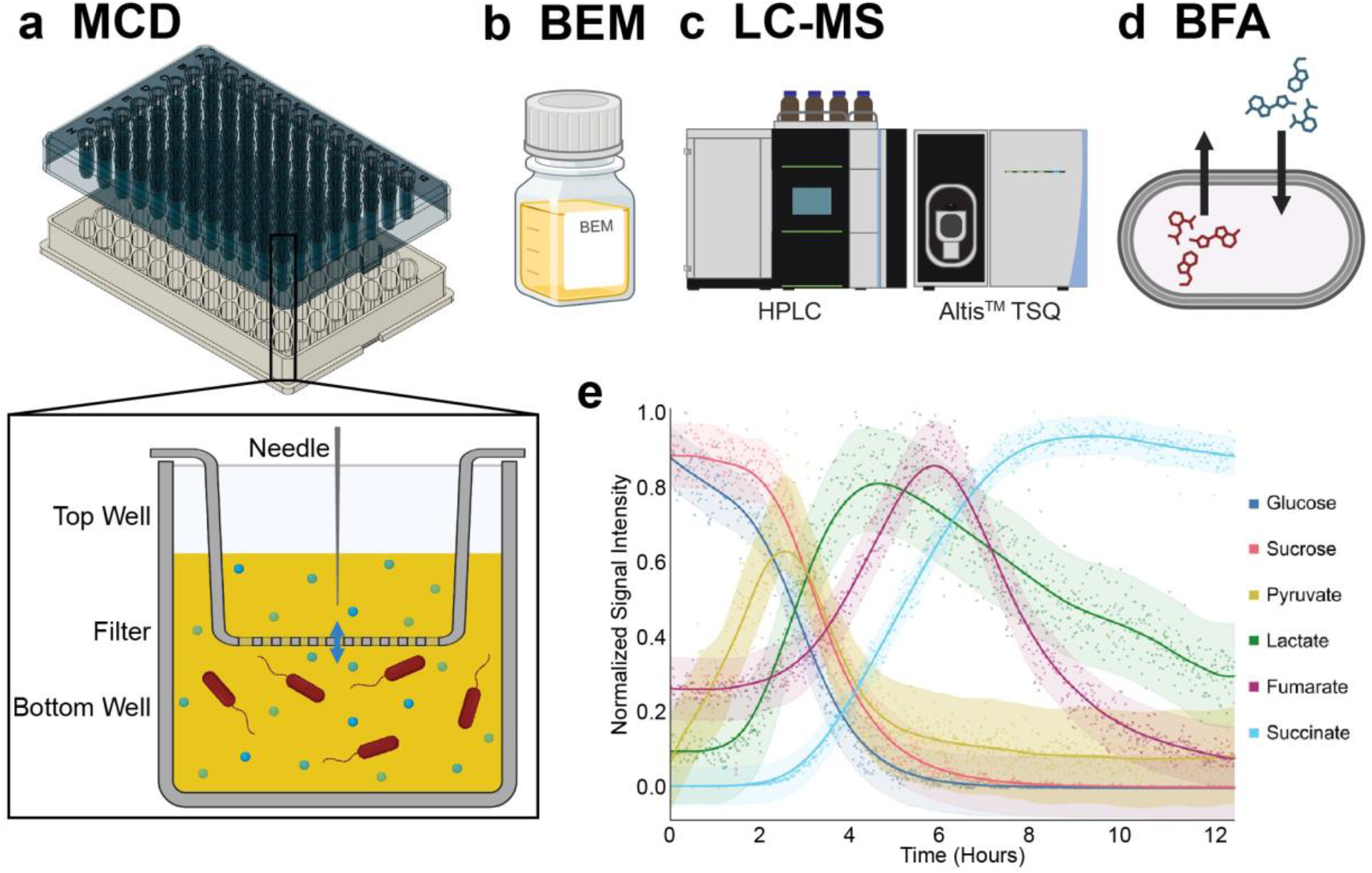
LC-MS system for time-resolved metabolomics. (a) Bacterial cultures are grown in a custom MCD, and (b) incubated in chemically defined BEM. Samples are then (c) incubated at 37 °C in the LC-MS autosampler, and samples are collected at 50-second intervals. (d) Changes in metabolite levels are used to compute boundary fluxes for each observed metabolite. (e) Proof of principle of our automated system was demonstrated with averaged, normalized levels of central carbon metabolites in *E. coli* cultures over 12 hours. Figure was made with BioRender.

We demonstrated TUNA using a Thermo Fisher TSQ Altis™ platform equipped with a Vanquish™ ultra-high-performance liquid chromatography system. Targeted metabolomics on this platform was enabled via selected reaction monitoring with specific transitions identified for 98 metabolites adapted from previously established methods^4,13,14^. The Vanquish™ supports sampling of individual 96-well plates at as little as 50-second intervals, which allows either sub-minute temporal samples to be collected (Fig. 1e) or up to 72 different culture conditions to be monitored within an hourly sampling schedule. One key design element of TUNA is our use of the MCD, which allows us to collect long time series data with minimal deterioration in signal performance due to instrument fouling (coefficient of variation between 3.2-7.9%) (Supplemental Fig. 2) and minimal technical variability between wells (Supplemental Fig. 3).

To demonstrate our automated TUNA when used for high density temporal analysis, we monitored *E. coli* cultures for over 12 hours with 50-second intervals (Fig. 1e). Briefly, cultures were seeded at 1 × 10^8^ CFU/mL in BEM^11^ inside the MCD^10^, and transferred to the 37 °C autosampler. Samples were then harvested at 50-second intervals over 12 hours. As expected, we observed *E. coli* catabolizing glucose and sucrose, and producing pyruvate, fumarate, lactate, and succinate (Fig. 1e). Moreover, our temporal data showed the switch between the production and consumption of multiple metabolites, including pyruvate, lactate, fumarate, and succinate at approximately two, four, five, and ten hours, respectively (Fig. 1e). These data illustrate TUNA’s ability to automatically capture temporal dynamics.

We then used chromatography enabled by TUNA to analyze the metabolic capacity of two closely related *Enterobacterales* species (*E. coli* and *Proteus mirabilis*) that commonly cause urinary tract infections. Strains (n=4 per species) were analyzed via a three-minute gradient, resulting in 144 time points over a seven-hour incubation (27-minute intervals) (Supplementary Fig. 4). As we reported previously^9,15^, isolates of the same species showed a high degree of metabolic homogeneity in their boundary fluxes (Supplementary Fig. 5a), but showed substantial interspecific differences (Fig. 2).

**Figure 2.**
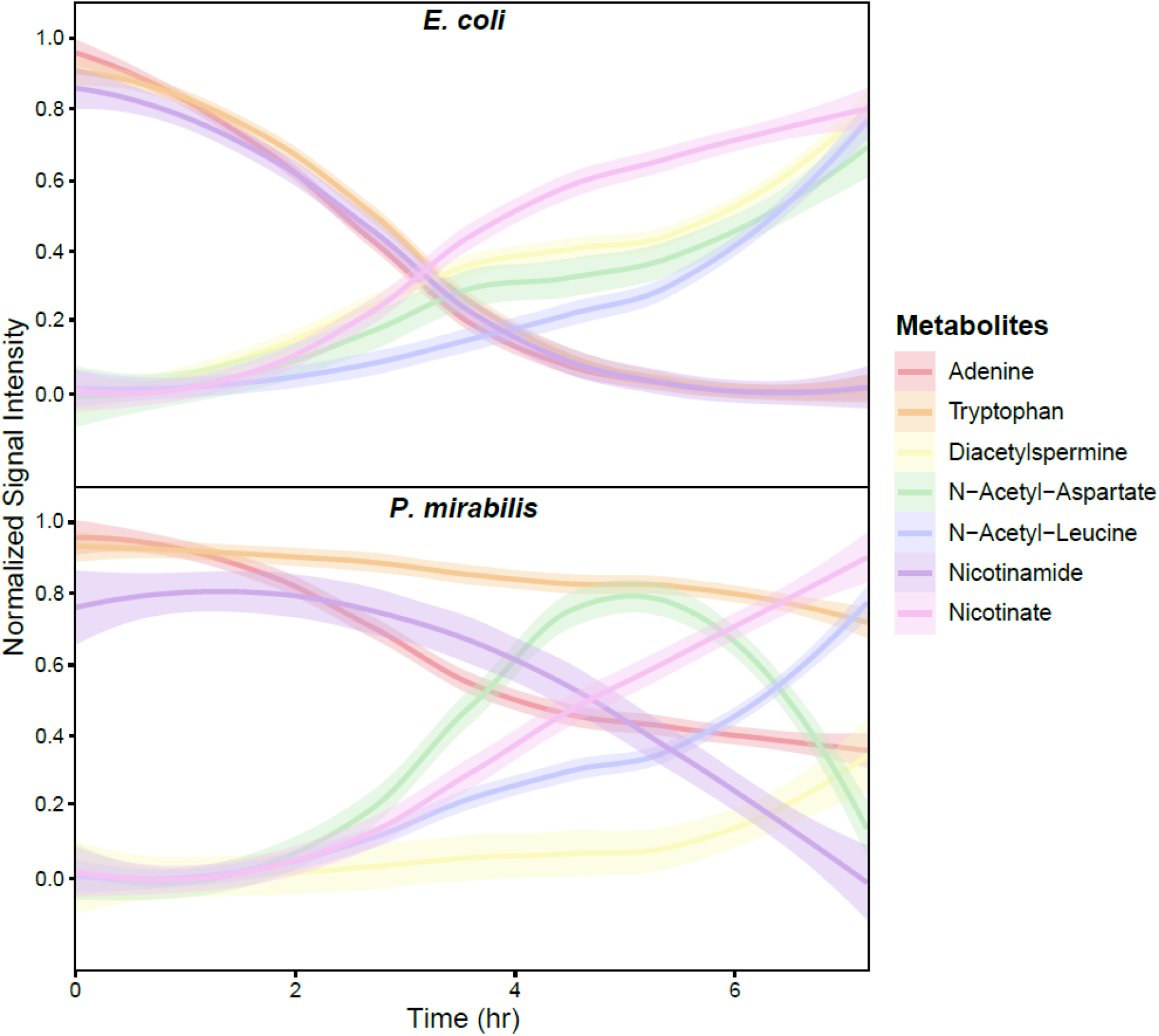
Select temporal metabolic phenotypes distinguishing *E. coli* and *P. mirabilis*. *E. coli* (n=4) and *P. mirabilis* (n=4) strains were grown at 1 × 10^8^ CFU/mL in BEM in the MCD. Signal intensities of metabolites were normalized by the maximum intensity and averaged amongst strains. The shaded area around each curve represents the 95% confidence interval, as generated by R using loess regression.

Moreover, we identified multiple novel temporal metabolic phenotypes that distinguished *E. coli* and *P. mirabilis* (Fig. 2). For example, *E. coli* produced N-acetyl-aspartate whereas *P. mirabilis* produced then consumed this molecule (Fig. 2). In addition, *E. coli* rapidly catabolized adenine, tryptophan, and nicotinamide, whereas *P. mirabilis* was slower to utilize these molecules (Fig. 2). Although metabolic profiles were generally stable within a species, we observed a few cases of metabolic heterogeneity between strains. For example, an *E. coli* strain (CFT073 ATCC 700928) was an outlier within *E. coli* due to its lack of ornithine consumption and elevated cadaverine production (Supplementary Fig. 5b). Interestingly, ornithine and cadaverine metabolism in *E. coli* have been previously linked to acid resistance^16^ and decreased antibiotic susceptibility^17^. Collectively, these observations show that TUNA can capture complex temporal metabolic dynamics that differentiate species and strains.

In summary, we introduce TUNA, an automated sampling system that can monitor the metabolic dynamics of cultures over time under physiological conditions. TUNA can be implemented in routine laboratory settings, provides physiologically relevant data, and allows a single condition to be monitored at 50-second intervals or up to 72 separate conditions to be monitored at hourly intervals. The key distinctions of TUNA from the solutions proposed previously include: (1) the ability to capture a broad transect of metabolites; (2) compatibility with chromatography, which enables robust metabolite assignments and minimizes ion suppression; (3) it does not inject live cells directly into the MS, and thereby minimizes fouling of the source and ion optics. Most importantly, TUNA is capable of capturing subtle temporal dynamics between multiple cell conditions that are analyzed in parallel. TUNA has several obvious applications, including mapping hierarchical nutrient utilization of cells^1^, monitoring drug-induced perturbations in metabolism^18^, and capturing ephemeral phenotypes that may not be present at all time points. Despite these advantages, some limitations in our system should be considered. Since cultures are grown in the autosampler, cells must be compatible with atmospheric gas conditions and steps must be taken to minimize evaporation (we added agarose gel to the reservoirs outside the MCD sampling wells). Provided that these limitations are addressed, TUNA can automatically capture time-resolved metabolomics data under physiologically relevant conditions using conventional laboratory instrumentation.

## Acknowledgements

This work was funded by the Natural Sciences and Engineering Research Council (NSERC) [DG 04547] supported by the Research Excellence Chair. Data were acquired at the Calgary Metabolomics Research Facility (CMRF) at the Alberta Centre for Advanced Diagnostics, which is supported by PrairiesCan (000022734) and the Canada Foundation for Innovation (CFI-JELF 34986). This work was made possible in part by a research collaboration agreement with Thermo Fisher Scientific.

## Author Contributions

R.A.G. designed and developed the temporal metabolomics monitoring platform. C.C.Y.C. performed experiments to test this new platform. C.C.Y.C. and I.A.L. wrote the manuscript.

## Methods

### Microbial culture preparation in the MCD

Four strains of *Escherichia coli* (MG1655 (ATCC 700926), CFT073 (ATCC 700928), ATCC BAA-2776, ATCC 11775) and four strains of *Proteus mirabilis* (ATCC 35659, ATCC 7002, ATCC BAA-2792, ATCC BAA-2791) were grown in either biomarker enrichment medium (BEM)^11^, or Roswell Park Memorial Institute (RPMI) 1640 medium (Thermo Fisher Scientific). These strains were first grown overnight from cryogenic stocks, and then overnight cultures were seeded at 0.34 McFarland (approximately 1 × 10^8^ CFU/mL) in fresh media. Density-normalized cultures were transferred into an in-house custom 96-well plate, the microbial containment device (MCD), consisting of the top wells (120 μl) overlaying the bottom wells (250 μl), separated by a 0.2 μm filter.^10^ To mitigate evaporation, only the inner wells were used and the outer trench surrounding the wells were filled with 1% agarose gel prior to the experiment. Technical replicates of each density-normalized culture were loaded in a column of bottom wells in the MCD (250 μl per well). Additional medium was loaded into the top wells (110 μl per well). Then, the MCD was placed inside the autosampler of the liquid chromatography-mass spectrometry (LC-MS) instrument set to an internal temperature of 37 °C.

### Automatic sampling process

Incubation duration was tracked starting from the time the MCD was loaded into the LC-MS autosampler. With the incorporation of the newly developed method described in the following sections, the LC-MS instrument can analyze samples at a rate of three minutes per culture. The autosampler first injected 1 μL from one single replicate of each culture condition. Then, another round of replicates was sampled, continuing until all wells were analyzed once before returning to the first replicates that were sampled. The sampling was allowed to continue for 24 hours.

### Chromatographic parameters

Sample incubation and analysis were performed on a Vanquish™ Flex Ultra-High-Performance Liquid Chromatography (UHPLC) System (Thermo Scientific). Chromatographic separation of targeted metabolites was achieved using a binary solvent mixture of 20 mM ammonium formate at pH 3.0 in LC-MS-grade water (Solvent A) and 0.1% (v/v) formic acid in LC-MS-grade acetonitrile (Solvent B) in conjunction with a 50 × 2.1 mm Syncronis™ Hydrophilic Interaction Chromatography (HILIC) Column with a 2.1 μm particle size (Thermo Scientific). Using this liquid chromatography system, the following solvent gradient was used at a flow rate of 1.0 mL/min: 0–0.2 min, 95% B; 0.2–0.7 min, 95–5% B; 0.7–0.95 min, 5% B; 0.95–1.2 min, 5-95% B; 1.2–2 min, 95% B. The sample injection volume for all analyses was 1 μL and column compartment temperature was 30 °C.

### Mass spectrometry parameters and data analysis

Data were acquired on a TSQ Altis™ Plus Triple Quadrupole Mass Spectrometer (Thermo Scientific) using the following heated electrospray source parameters: spray voltage of +3000 V/-2500 V, sheath gas of 35 (arbitrary units), auxiliary gas of 15 (arbitrary units), sweep gas of 2 (arbitrary units), ion transfer tube temperature of 275 °C, and auxiliary gas temperature of 300 °C. Data acquisition was performed using selected reaction monitoring. The optimized parameters for 98 metabolite targets were attained via direct infusion of individual authentic standards obtained from Sigma-Aldrich and Fisher Scientific. All data were acquired with mixed scan mode using a dwell time of 1 ms, Q1 resolution of 0.7, and Q3 resolution of 1.2. Data acquisition was carried out using Xcalibur™ v4.2 software (Thermo Scientific) and all spectral data were integrated using El-MAVEN v0.12.0^19^.

## Notes

### Competing Interest Statement

The authors have declared no competing interest.

