## Supplementary Figures for "LC-MS system for automatically collecting time-resolved metabolomics data of cultured cells"

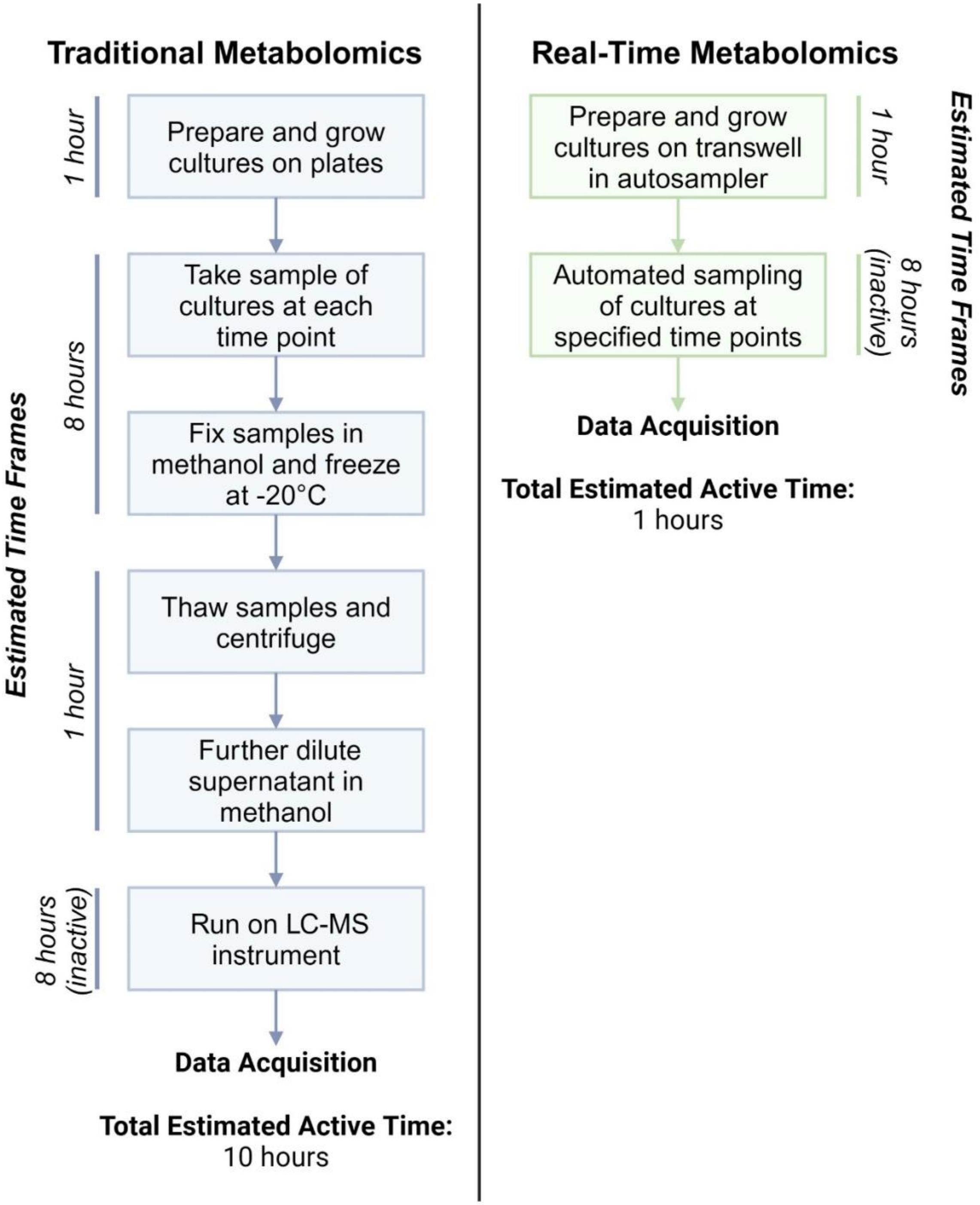


**Supplementary Figure 1. Experimental design of traditional metabolomics compared to our automated TUNA system.** The active and inactive times required for each step were estimated for an eight-hour bacterial growth course in which cultures were growth on a single 96-well plate. In traditional metabolomics experiments[3,4] (left), microbes are seeded in growth medium from cryostocks and grown overnight. The overnight cultures are then used to inoculate the experimental cultures to a normalized starting microbial density. After inoculation, the cultures are grown in the incubator, and at each time point, the cultures are manually sampled and fixed in methanol to halt further microbial metabolic activity. This sampling step may be repeated at multiple time points to acquire time-resolved metabolomics data. Culture samples can either be frozen and stored, or the samples can be further diluted in methanol and then analyzed on the LC-MS instrument. Whereas in the automated TUNA system (right), the preparation of the microbial cultures in the MCD is the only manual step required.

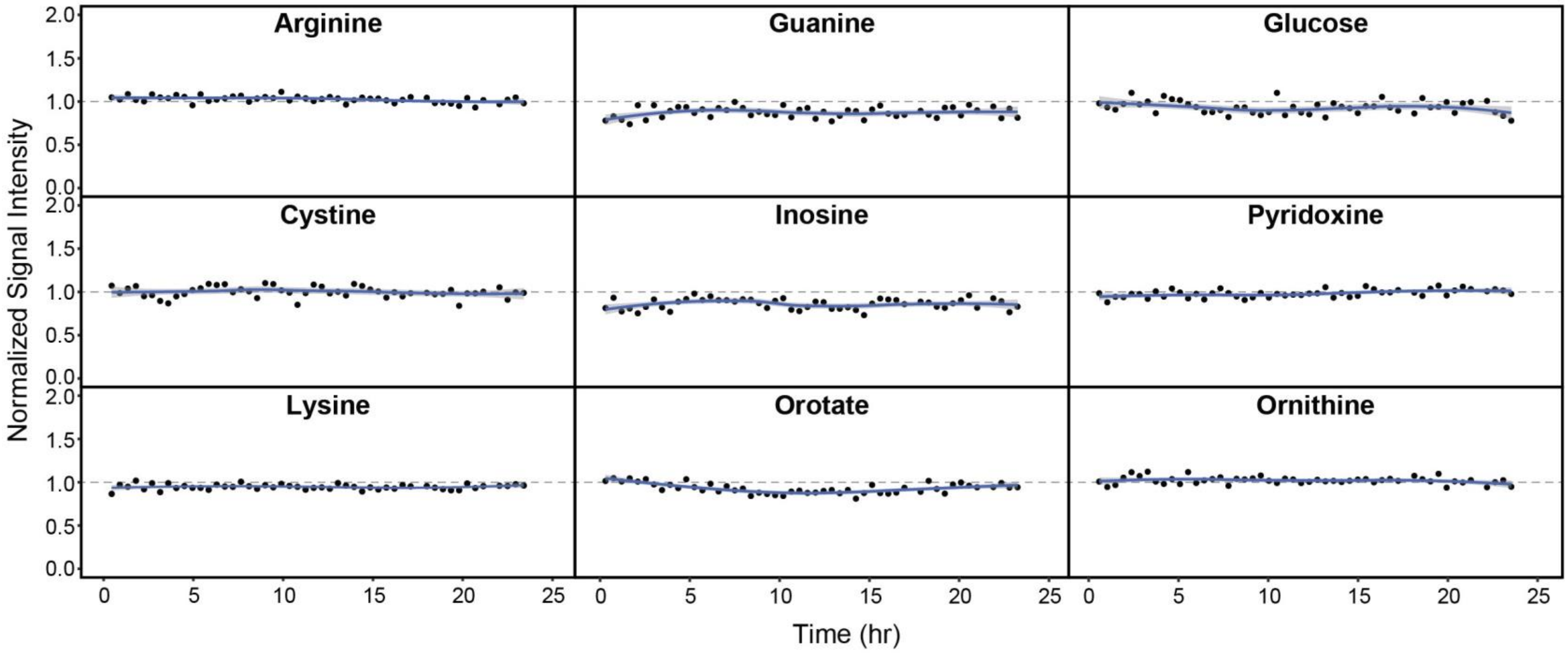


**Supplementary Figure 2. Demonstration of instrument performance stability over time.** Uninoculated BEM media was analyzed in the MCD over 24 hours at 27-minute intervals. The signal intensities of select metabolites were normalized against the starting signal intensity. The normalized signal intensities of arginine, cysteine, lysine, guanine, inosine, orotate, glucose, pyridoxine, and ornithine over time are used to demonstrate TUNA's performance stability. The grey shaded area around each curve represents the 95% confidence interval, as generated by R using loess regression. The coefficient of variation of these metabolites is between 3.2-7.9%.

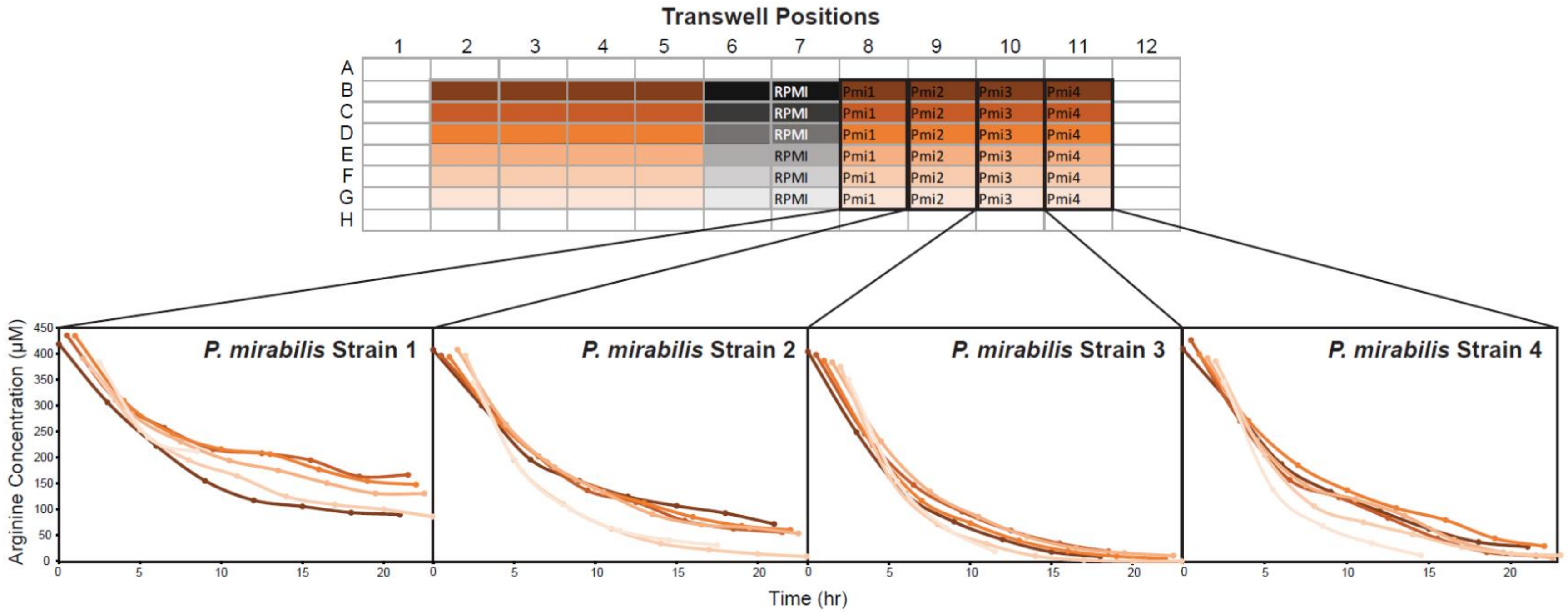


**Supplementary Figure 3. Well-to-well differences of arginine signal intensity in MCD.** To set up TUNA, six technical replicates originating from the same culture were positioned on the same column in the MCD, and only inner wells were utilized for the experiment. For example, replicates of *P. mirabilis* strain 1 were growth in B-8 to G-8, replicates of *P. mirabilis* strain 2 were growth in B-9 to G-9, and so forth. Graphs displayed differences in the signal intensity of arginine between the individual replicates of each strain when grown in RPMI media.

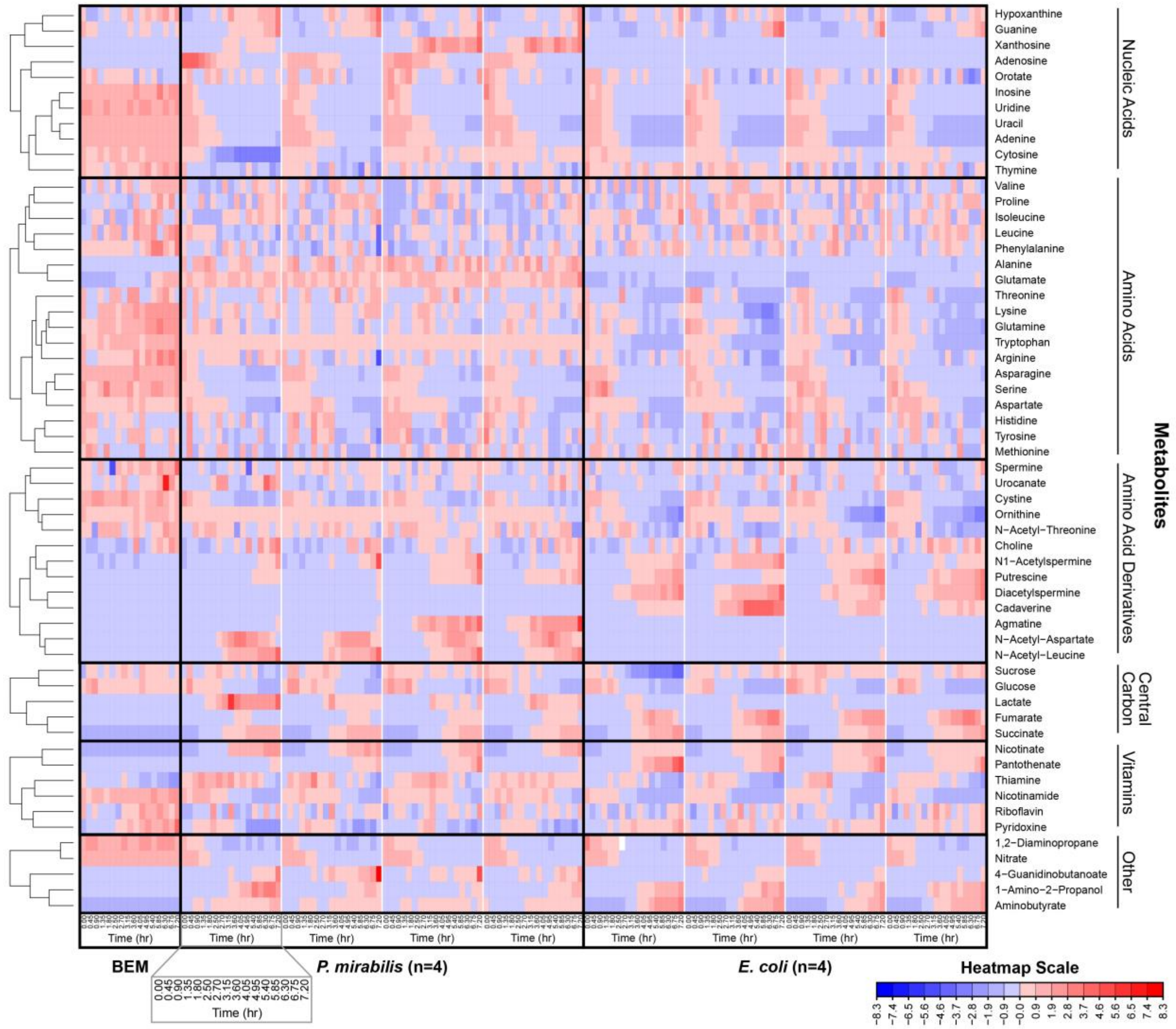


**Supplementary Figure 4. Heatmap of the signal intensities of common metabolites in *in vitro E. coli* and *P. mirabilis* cultures monitored with TUNA.** *E. coli* (n=4) and *P. mirabilis* (n=4) strains were grown at $1 \times 10^8$ CFU/mL in BEM media inside the MCD. The automated TUNA setup allowed each bacterial culture to undergo targeted LC-MS analysis every three minutes, resulting in 27-minute time points. Red indicates higher metabolite signal intensity and blue indicates lower metabolite signal intensity.

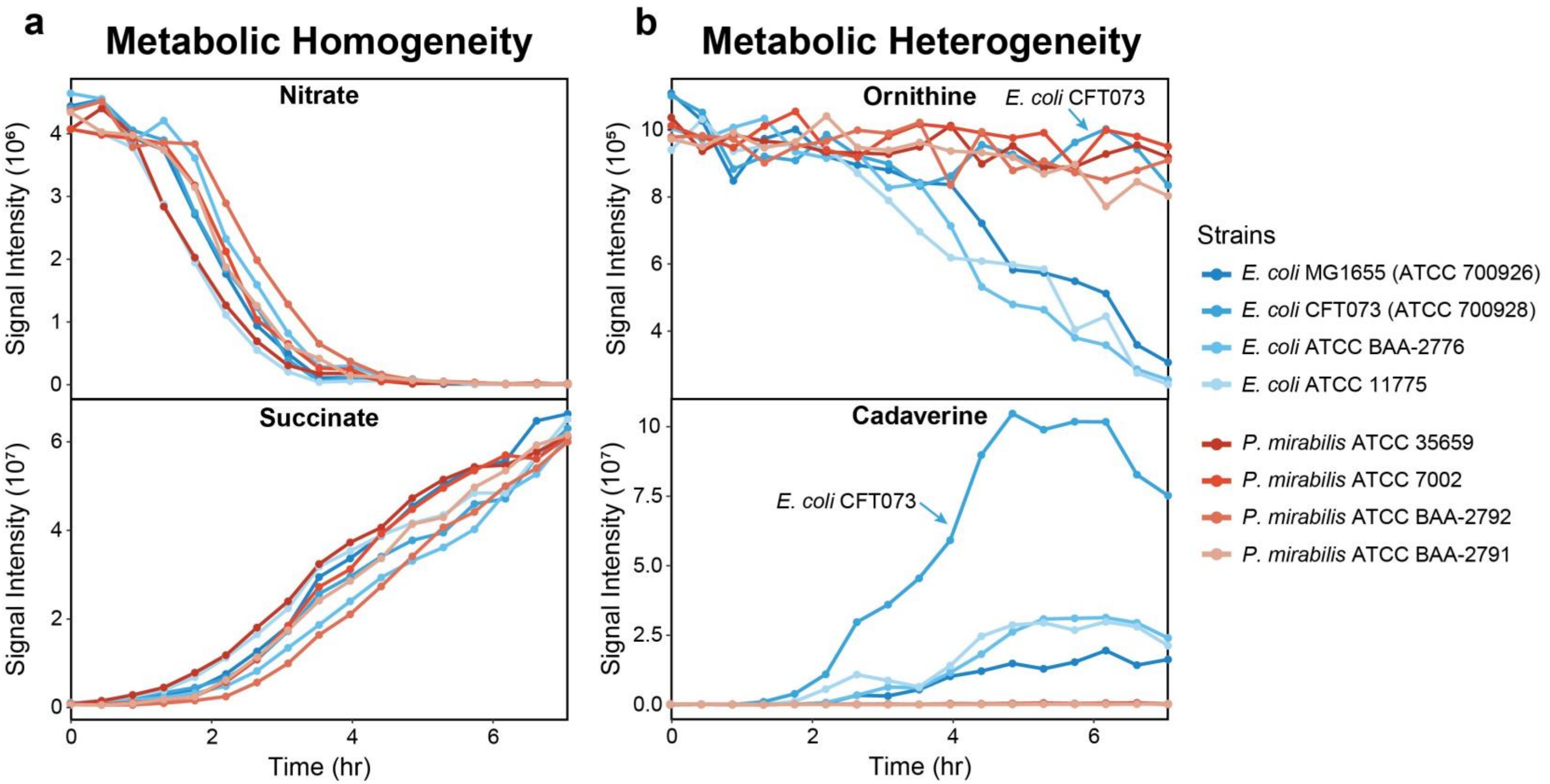


**Supplementary Figure 5. Signal intensities of select metabolites produced or consumed over time by individual *E. coli* and *P. mirabilis* strains.** The levels of cadaverine, nitrate, ornithine, and succinate in bacterial BEM cultures were monitored with TUNA for seven hours. These data demonstrate metabolic homogeneity (a) and heterogeneity (b). An *E. coli* strain (CFT073) demonstrated metabolic phenotypes distinct from other *E. coli*, as labeled in the graphs.
